# Efficient Assembly and Verification of ZFNs and TALENs for Modifying Porcine *ApoE* gene

**DOI:** 10.1101/013714

**Authors:** H R Xu, T Li, Y Guo, H F Li, L Wang, Z Y Zhang, X Wang

**Author notes:** Xu H R, College of Animal Science and Technology, Northwest A&F University, YangLing, Shaan’xi. Li T, College of Animal Science and Technology, Northwest A&F University, YangLing, Shaan’xi. Guo Y, College of Animal Science and Technology, Northwest A&F University, YangLing, Shaan’xi. Li H F, College of Animal Science and Technology, Northwest A&F University, YangLing, Shaan’xi. Wang L, College of Animal Science and Technology, Northwest A&F University, YangLing, Shaan’xi. Zhang Z Y, College of Animal Science and Technology, Northwest A&F University, YangLing, Shaan’xi. Corresponding author: Wang X, College of Animal Science and Technology, Northwest A&F University, YangLing, Shaanxi. Corresponding author: Wang X, TaiCheng Road 3, College of Animal Science & Technology, Northwest A&F University, YangLing, Shaan’xi, P. R. China.

## Abstract

Zinc-finger nucleases (ZFNs) and transcription activator-like effector nucleases (TALENs) are powerful tools for genome engineering. These synthetic nucleases are assembled with programmable, sequence-specific DNA-binding domain and a non-specific FokI cleavage domain. Apolipoprotein E (*ApoE*) gene polymorphism is associated with cardiovascular outcomes, including ischaemic stroke and coronary heart disease (CHD). So the objective of this study is to create mutations of *AopE* gene by ZFNs and TALENs technology. Here, we used the Context-dependent assembly (CoDA) method to design and screen ZFNs specifically targeting with *ApoE* gene. The targeted cleavage capacity of these ZFNs was validated in yeast system and HEK 293T cells. Meanwhile, an efficient assembled TALENs to target *ApoE* gene in HEK 293T cells was as a control. The results showed that both ZFNs and TALENs worked on *ApoE* gene with similar high-efficiency cleavage capability. The result would provide efficient methods for genome editing, so as to get disease model for gene therapy for the further study.

## Introduction

Synthetic zinc finger nucleases (ZFNs) and transcription activator-like effector nucleases (TALENs) are widely used in genomic integration, and formed by fusing programmable specific DNA binding module to the non-specific nuclease domain of FokI endonuclease (Cathomen and Joung 2008). The construct of ZFNs and TALENs can create a double-strand break (DSB) to either stimulate intracellular DNA repair by homologous recombination (HR) with donor DNA or induce gene mutation by non-homologous end joining (NHEJ) when without a donor DNA (Jeggo 1998; van Gent *et al*. 2001). ZFNs and TALENs-mediated genome engineering has been used in many organisms, including cultured cells, as well as in live animals and plants (Kim and Kim 2014). Apolipoprotein *E (ApoE)* is an important component of plasma apolipoprotein, comprised by 299 amino acids. *ApoE* gene has three alleles, named E2 and E3 and E4. Due to mutation in 112^th^ of *ApoE*4, Cys at this site was substituted for Arg. This mutation in *ApoE*4 has significant influence on coronary heart disease, cerebral infarction and senile dementia etc. diseases (Scarmeas *et al*. 2002).

Considering together, we sought to create mutations in *ApoE*4 gene by using efficient genome editing methods of ZFNs and TALENs. Therefore, taking porcine *ApoE* gene as the target site, two specific ZFNs were generated by utilizing CoDA assembly method and one TALENs was assembled by our simple method in this study, which would provide efficient methods for genome editing and lay foundations for producing *ApoE* gene defected cell lines, so as to establish *ApoE* gene defected disease model for gene therapy for further study.

## Material and Methods

### Zinc Finger Targeter Selection

The two ZFNs targeting porcine *ApoE* gene was designed by the ZiFiT website (http://zifit.partners.org/ZiFiT/ChoiceMenu.aspx), which allows users to copy and paste their sequences of interest and returns the available CoDA ZFN sites as well as the DNA sequences that will encode a particular array.

### Oligonucleotide Design and Synthesis

For this study, oligos were designed based on the target sequence, and were synthesized by Biosune Lt. Company (Shanghai). The primer pairs used for plasmids construction, detection and sequencing in this work are presented in Table 1.

### Overlapping PCR to Obtain the ZFP Fragments

The ZFP fragments of *ApoE* (ApoE-ZFPs) were amplified by PCR and the products were purified using the Gel Extraction Kit (Vigorous, Beijing). The total reaction volumes were as follows: 10 × PCR Buffer 5.0 μL, dNTPs 4.0 μL, forward and reverse primer 1.0μL, pfu (5 U/μL) 0.5 μL, DNA 1 μL, adjusting ddH_2_O to 50 μL. The reaction protocol was: 95° for 5 min, 5 cycles of: 94° for 30 sec, 52° for 30 sec, and 72° for 30 sec, then add corresponding primers, followed by 30 cycles of: 94° for 30 sec, 52° for 30 sec, 72° for 30 sec and 72° extension for 5 min.

### Construction of ZFN Expression and Reporter Vectors for Yeast-two-hybrid System

The left fragment of ZFP (LZFP) was digested by *Xba*I and *Bam*HI (NEB), while the right fragment of ZFP (RZFP) was digested by *Not*I and *Bam*HI. Then the LZFP and RZFP were cloned into *Xba*I-*Bam*HI site of JMB440-FokI backbone and *Not*I-*Bam*HI site of JMB405-FokI vector, respectively. Plasmid DNA was isolated with the E.Z.N.A. Plasmid Mini Kit (OMEGA) and sequenced to confirm whether the left and right finger inserted into the colonies or not. These vectors were named as LZFN-JMB440 and RZFN-JMB405. For the corresponding reporter vector, the oligonucleotides of Ap-BS1, Ap-BS2 (Table 1) were annealed directly to obtain the double strand DNA with *Not*I-*Bam*HI site and cloned into JMB52 vector, named as ZFBS1-JMB52 and ZFBS2-JMB52.

### Screening Specific ZFNs in Yeast-two-hybrid System

To test whether ZFNs work or not, we co-transform the plasmids of LZFN-JMB440, RZFN-JMB405 and reporter vectors ZFBS-JMB52 into yeast AH109 (Figure 1). The transformed products were cultured in SD solid medium (-His-Ade-Trp-Leu). As control, we transformed reporter plasmid only and cultured in SD solid medium (-His-Ade+Trp+Leu). After 3-day’s culture at 30°, reporter plasmids were recovered from plates (-His-Ade-Trp-Leu) and control plates, respectively. The specific fragments were amplified by primer FADH1 and gal4ADR. The primer sequences were listed e in Table 1.

**Figure 1.**
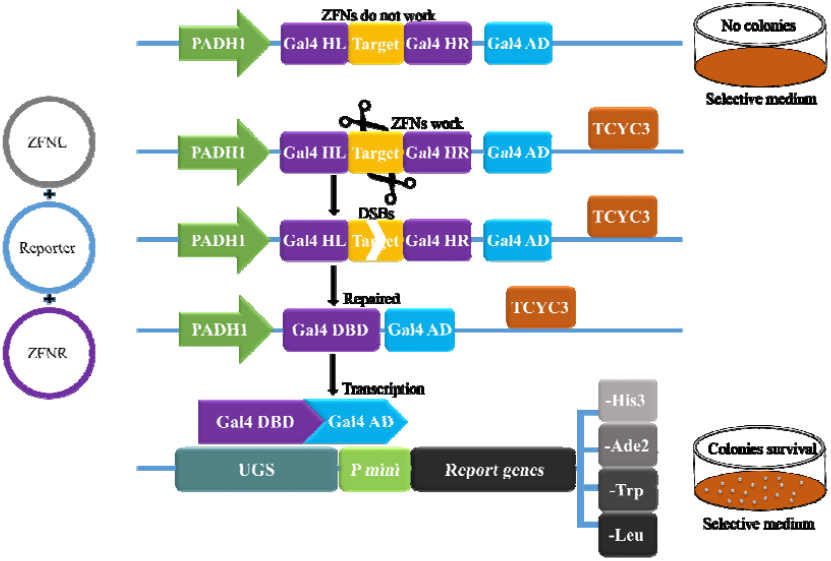
Diagram describing yeast-based ZFNs screening and validation system. ZFNs and reporter co-transform into yeasts. If ZFNs do not work, no colonies can survival with selective medium. With this validation system, we can screen the working ZFNs and with control we can obtain a sketchy efficiency of ZFNs by counting colonies number.

### Engineering ZFNs Eukaryotic Expression and Reporter Plasmid for Mammalian Cells

The aforementioned LZFP and RZFP were cloned into PST1374-L vector. The plasmids (PST1374-ZFNL, PST1374-ZFNR) were extracted and sequenced. The protocol of constructing reporter vector for mammalian system (Figure 2) was the same as that for yeast-two-hybrid assay. The oligonucleotides of BS1 and BS2 were annealed, then inserted into psdRED backbone. The principle of this reporter verification system was the same as Hyojinkim described (2011).

**Figure 2.**
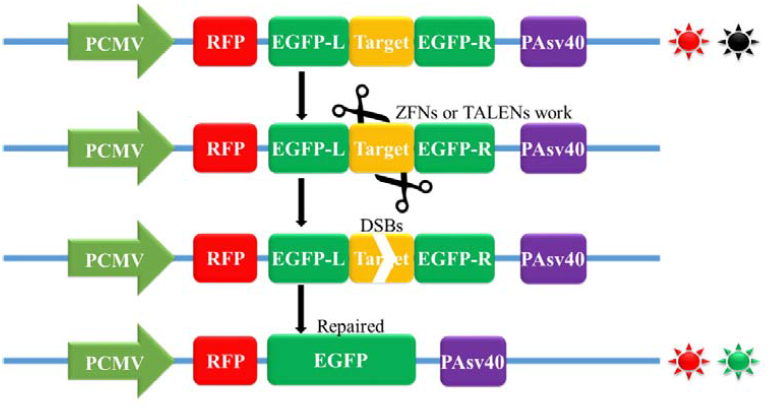
Red-Green light report system. In HEK 293 cells, ZFNs and TALENs cut target sequences to induce EGFP repare. In the green field we can obtein green spots if they work. In the same way, we obtein effiencies by counting green/red spots.

### Validate Activity of ZFNs in mammalian cells

HEK 293T cells were maintained in DMEM contained 10% fetal bovine serum and seeded into 6-well plates before transfection. A mixed plasmids comprising ZFNL-pst1374 construct, ZFNR-pst1374 construct and BS-psdRED vector were co-transfected in 3.2 × 10^5^ cells per well using lipofectamine2000 following the manufacturer’s instruction (Invitrogen). As control, cells were transfected with BS-psdRED vector only. After cultivating for 36 h at 37°, the fluorescence was observed by fluorescence microscope system and then DNA was extracted 12 h later and detected by PCR.

### TALE Targeter Selection and Synthesis

TAL Effector Nucleotide Targeter 2.0 was used to find porcine *ApoE* gene targeter (https://tale-nt.cac.cornell.edu/node/add/talen). We chose one target site for assembling TALENs based on our construction protocol of TALE (Zhang et al., 2013). The primers used for plasmids construction, detection and sequencing were synthesized by Shanghai Biosune Company. The entire primer information was presented in Table 1.

### Construction of TALENs Eukaryotic Expression Vectors

Three tetramer plasmids chosen for construction of TALENs were amplified with primers Tet-Fv\Tet-R1, Tet-F2\Tet-R2 and Tet-F3\Tet-Rv, respectively. Primer sequences were presented in Table 3. Typically, the PCR reaction was: 20 ng plasmids, 1 uL of each forward and reverse primer (5 uM), 0.5 uL Taq DNA polymerase (5 U/uL), 5 uL of 10 × Taq polymerase buffer, 5 uL of 10 mM dNTP, finally add ddH_2_O to 50 μL. PCR protocol was 94° for 5 min; 94° for 30 sec, 57° for 30 sec, 72° for 30 sec, 35 cycles; 72° for 5 min. The amplified tetramer was 465 bp, and purified from gels with PCR production purification kit. The three tetramers (50 ng each) and pST-TALEN-Backbone (150 ng) were added into a 10 uL volume of cut/ligation reaction mix with 1 uL of *Bsm*BI(10 U/uL), 1 uL of T4 DNA ligase (400 U/uL), 1 uL of 10 × T4 DNA ligase buffer and ddH2O. The cut/ligation reaction was done in a thermocycler with 42° for 5 min, 16° for 5 min, 30 cycles. Then transformation was carried out with DH5α using 5 uL cut/ligation reaction mix. The transformation mix was plated on LB solid medium containing 100 ug/mL ampicillin and incubated at 37° for 12 h. The plasmids were obtained with Plasmid Mini Kit, then verified by *Xba*I/*Bam*HI digestion, then the verified plasmids were sequenced with primers Seq-F and Seq-R. The completely right plasmid was named by pST-ApoE-TALEN.

### Engineering TALENs Reporter Plasmid for Mammalian Cells

For the corresponding reporter vector, the oligonucleotides of Ap-BS (Table 1) were annealed directly to obtain the double strand DNA with *Not*I-*Bam*HI site and cloned into psdRED backbone vector. The principle of this reporter verification system was the same as Hyojinkim described (2011).

### Validate Activity of TALENs in Mammalian Cells

HEK 293T cells were cultured with DMEM contained 10% fetal bovine serum and seeded into 6-well plates before transfection. The transfection was the same as above described.

## Results

### Three Zinc Finger Sequence Acquired and Oligonucleotide Design

Two pairs of ZFNs target sites were chosen from porcine *ApoE* gene near the mutation site by utilizing CoDA platform.

BS1: 5’-<u>*GGCGGCGCA*</u>ggccgcc<u>*GTGGGCGCC*</u>-3’;

BS2: 5’-<u>*GACCACCGA*</u>ggagc<u>*TGCGGAGCC*</u>-3’.

In CoDA database, specific three zinc finger proteins were screened, which recognized 9 bp of left and right, respectively, following the method as described by Jeffry D Sander et al (2010) (Table 1). After ZFP amino acid being translated, the primers were designed to amplify the segment of ZFP (Table 2).

### Engineering ZFNs Activity Assaying

A 270 bp fragment of ZFP was got by overlap PCR (Fig. 3A). The selected ZFPs were successfully cloned into a ZFN expression vector. Candidate ZFN constructs were assayed for specific cleavage activity in yeast-two-hybrid (Y2H) system as previously described (Fig. 3B). The Y2H reporter assay revealed that these two ZFNs constructs both have high efficiency for reporter gene modification in yeast (>90%). Based on this result, the cleavage activities of our selected ZFNs were carried out in mammalian cells. The capabilities were assayed by repairing the reporter gene via homologous recombination. For both two ZFNs target sites, eukaryotic expression plasmids and corresponding reporter vectors were constructed (Fig. 3C), then transfected into HEK 293T cells. After transfection about 48 h, the results showed that these two pairs of ZFNs exhibited a substantially higher mutation frequency (~10%-20%) (Figure 4).

**Figure 3.**
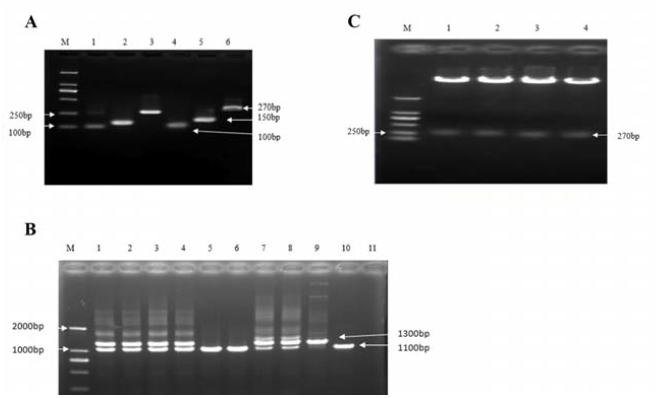
Assembling ZFNs. A, The PCR products of ZFPs; B, The PCR products of Gal4BD gene. We detected 1100 bp stripes to identify whether ZFNs work. No.9 is negative control, No.10 is positive control; C, The digestion analysis results of ZFNs eukaryotic expression vectors (M: DL2000 DNA Marker)

**Figure 4.**
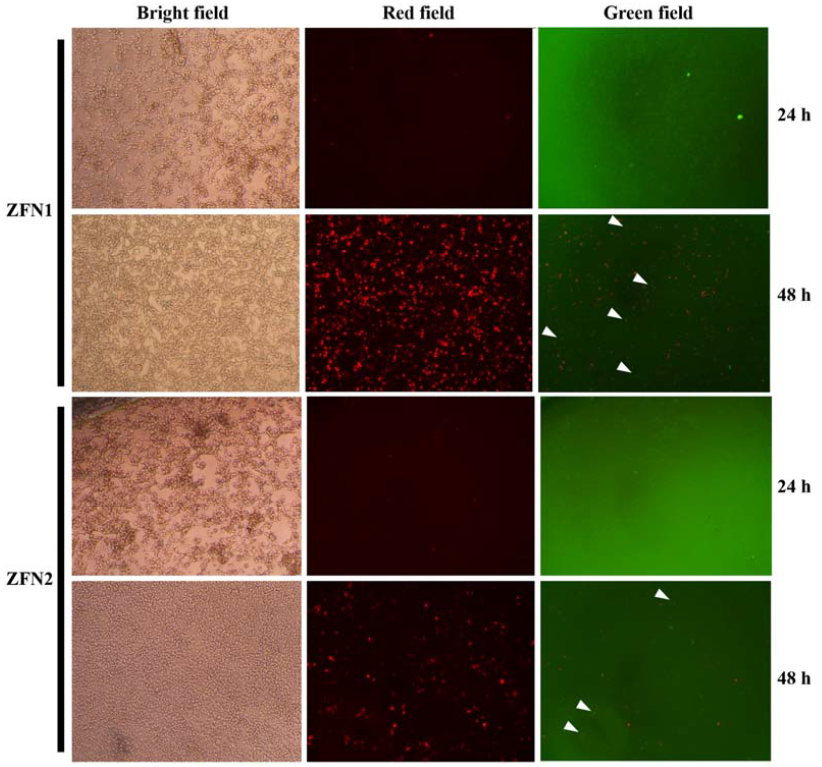
Fluorescence results treated by ZFNs in HEK 293T cells. ZFN1 and ZFN2 respectively co-transfect into HEK 293 cells with reporter. If ZFNs work, we can obtain green spots (white triangles) about 48 h later. In the same way, we can obtain ZFNs efficiency through counting green (sometimes a bit saffron) spots and red spots. ZFN1 is over 20%, while ZFN2 is about 10%.

## Engineering TALENs and Assaying the Activity

### Left and Right TALE Tetramers Chosen for *ApoE* Gene

The TALENs target binding site designed by online tools TAL Effector Nucleotide Targeter 2.0 was as follows:

5’<u>*GGGCGCCGACAT*</u>ggaggacgtgcgcaacc<u>*GCTTGGTGCTCT*</u>-3’. The tetramers for the construction of left and right TALE were in table 3.

The tetramers were assembled by our simple and efficient method (Fig. 5A), then the specific TALENs was constructed and detected (Fig. 5B). The TALENs were co-transfected with reporter vector into HEK 293 cells. After 48 h, green light spots were obtained in the working group (Figure 6), while the control group without TALENs only showed in red light.

**Figure 5.**
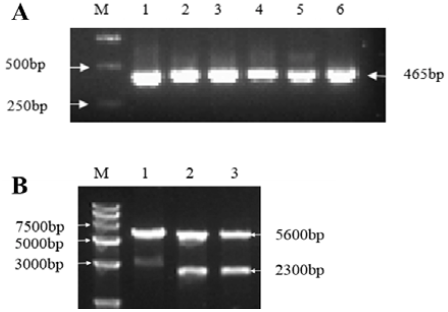
Assembling TALENs. A, Amplified tetramers 1-6 by PCR; B, Restriction detection of pST1374-TALEN-*ApoE*

**Figure 6.**
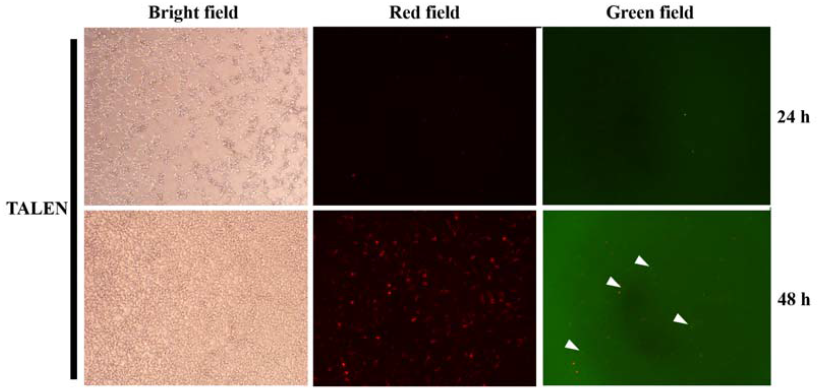
Fluorescence results treated by TALENs in HEK 293T cells. Under green field, we observed green spots (white triangles) about 48 h later and estimated TALENs efficiency is about 15%.

## Discussion

Until now, as a new genome modification tool, ZFN has been used widely in a variety of research fields, especially in gene therapy and clinical application. Here, we rapidly assembled ZFN arrays into a plasmid vector containing heterodimeric FokI nuclease domains using CoDA method at a comparatively low cost and time-consuming experimental selection. The activity of ZFNs was tested in yeast system preliminary (Figure 1) and then in HEK 293T cells (Figure 2). The results indicated that repair efficiency of our assembled ZFNs was as high as 95% in yeast, and about 20% in HEK 293T cells. These results fully explained CoDA method is an inexpensive, rapid assembly procedure to screen zinc finger protein.

Late-onset Alzheimer’s disease (LOAD) risk is strongly influenced by genetic factors such as the presence of apolipoprotein Eε4 allele (referred to here as *APOE*4),as well as non-genetic determinants including ageing. These data implicate an *APOE*4 associated molecular pathway that promotes LOAD (Rhinn *et al*. 2013). Recently one meta-analysis suggests that at least one *ApoE*ε4 allele has higher risk suffering AD than controls in Chinese population (Liu *et al*. 2014).

How to reveal *ApoE* gene function to obtain functionally and clinically relevant information means a lot for curing a series of human diseases, such as AD. A major issue for AD research is lack of animal models that accurately replicates the human disease, thus making it difficult to investigate potential risk factors for AD such as head injury (Bates *et al*. 2014). Currently, the main animal model for *ApoE* gene study is mice (de Castro *et al*. 2014; Klein *et al*. 2014).

While due to the high similarities in anatomy, genetics and pathophysiology with humans, pig has been a top choice as human health and disease model (Walters *et al*. 2012). Human late-onset diseases such as Parkinson’s disease (LRRK2, SNCA) and Alzheimer’s disease (TUBD1, BLMH, CEP192, PLAU) may also occur in pigs (Groenen *et al*. 2012). In this study, we successively constructed both ZFNs and TALENs to target pig *ApoE* gene in eukaryotic cells. Efficient and convenient tools to knockout *ApoE* gene can provide technical support to produce biomedical model for human disease.

Since there are a variety of methods for study the function of genes, such as gene knockout, gene silencing, gene overexpression etc. Targeted gene knockdown by RNAi has provided a rapid, inexpensive, and high-throughput alternative to homologous recombination (McManus and Sharp 2002). However, knockdown by RNAi has unpredictable off-target effects, and provides only temporary inhibition of gene function. These restrictions impede researchers to associate phenotype with genotype directly and limit the practical application of RNAi technology (Gaj *et al*. 2013). ZFNs and TALENs are relatively mature technologies. These synthetic nucleases contain programmable DNA-binding domains and non-specific DNA cutting domains. This combination of simplicity and flexibility has catapulted ZFNs and TALENs to the forefront of genetic engineering (Gaj *et al*. 2013).

We used both ZFNs and TALENs to target *ApoE* gene, and their activities were detected in yeast and HEK 293 cells through red-green light report system. The results showed that ZFNs worked better than TALENs. Two pairs of ZFNs were constructed with CoDA methods. At first we detected their activities in JM109 cells. It’s easy to find out whether they can work or not with gald4 report system in the yeast (Wang *et al*. 2013). Then we co-transfect ZFNs expression vectors and report vectors into HEK 239 cells to detect if they could work in mammalian cells. The green spots were obtained in the treatment group, while the control group only showed red light without ZFNs transfection. ZFP is the core part of ZFNs which can decide whether ZFNs work or not. How to obtain specific zinc finger protein and detect its DNA-binding activity is the key of ZFN technology. By now mainstream methods of assemble ZFP are module assembly, OPEN and CoDA. Module assembly is simple and convenient, but it doesn’t consider the module context effect, so this method has low efficiency (Ramirez *et al*. 2008). Based on bacterial two-hybrid method, Joung’s group designed OPEN method for assembling ZFP (Maeder *et al*. 2008). This method based on a large ZFP library to fix module context effect. Although with OPEN method researchers have obtained high activity ZFP and improved the gene targeting efficiency, this solution is too complex to popularize. Joung’s team obviously realized this issue. In 2011, they created the CoDA protocol. Now with the OPEN ZFP information foundation, this protocol is feasible for most common organization. Through designating the middle ZF2 which characterizes universality, ZF1 and ZF3 are easy to be obtained. Although we tried both OPEN and CoDA methods to assemble ZFNs for targeting *ApoE* gene, the former solution took a long time to carry out. Meanwhile ZFNs’ efficiencies obtained with OPEN method were similar to these obtained with CoDA method.

Move over ZFNs? TALEN technology now has been brought into biotechnology. So far most of reports claimed that with TALENs it is easier to mutate or reform the target gene. We built two pairs of TALENs for targeting *ApoE* gene at the same time to test which kind of tools work better. Only considering the time for assembling ZFNs and TALENs, it is true that TALENs prevail ZFNs. We assembled TALENs only within one week (Zhang *et al*. 2013), while the assembling of ZFNs took about one and a half month. With the same red-green light system, we co-transformed TALENs’ expression vectors and report vectors into HEK 293 cells. About 24 hours later, we could detect the green light in the treatment group. 48 hours later, the pictures obtained with fluorescence inverse microscope showed that about 10% cells with green light spot. Three tetramers were used to assemble each side twelve monomers of TALENs for targeting *ApoE* gene. The binding sites were selected with the online tool: TAL Effector-Nucleotide Targeter 2.0 (Doyle *et al*. 2012).

The contiguous repeat of DNA-binding modules of TALENs affects the efficiency of targeting gene. TALE-repeats vary in numbers from 15.5 to 19.5 among most of naturally occurring TALE (Boch and Bonas 2010). Theoretically, the larger TALENs with more tetramers should have higher efficiency. But the fact is not like that. RVD composition as well as the size of the repeat array affects target specificity (Morbitzer *et al*. 2011). The in silico analysis revealed that TALE-repeats longer than 12 bp reached a plateau in TALEN-pair–binding specificity in the euchromatic regions of the genome (Katsuyama *et al*. 2013), so we chose 12 bp TALE-repeat in this study.

Assembling TALENs is much easier than ZFNs, but efficiency of targeting gene is not so remarkable. Typically TALENs protein size is bigger than ZFNs and TAL effectors are naturally occurring proteins from the plant pathogenic bacteria genus Xanthomonas. The Cys2-His2 zinc-finger domain represents the most common DNA-binding motif in eukaryotes and the second most frequently encoded protein domain in the human genome (Gaj *et al*. 2013). In consideration of immunogenicity, ZFNs which are ubiquitous in nature may not be an issue. But TALENs may have some problems in gene therapy. Clearly, how to deliver programmable nucleases to cells safely and effectively also affects the targeting efficiency.

Currently genetically engineered pig models are being used for analysis of gene function in various human diseases, development of new therapeutic strategies as well as production of biopharmaceutical products (Walters *et al*. 2012). We built both ZFNs and TALENs to target porcine *ApoE* gene and detected activities in HEK 239 cells with red-green light system. The preliminary application results show that this approach is effective and feasible.

## Acknowledgments

This research is supported by Scientific and Technological Project in Shaanxi province (No. 2014K02-07-01).

---

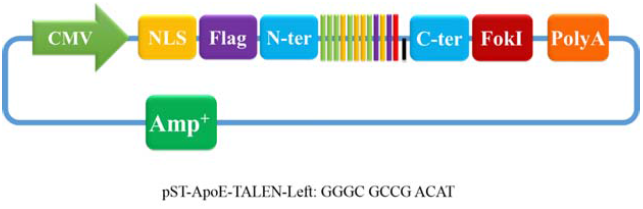

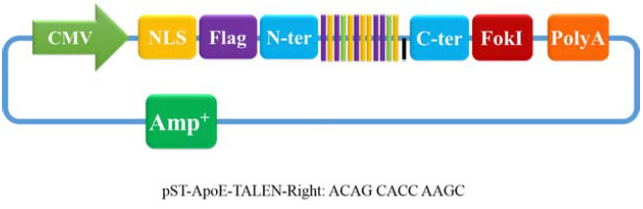

